# MEX3B is a positive pan-inflammasome regulator

**DOI:** 10.64898/2026.03.30.714824

**Authors:** Jason G. Cahoon, Tingting Geng, Duomeng Yang, Conner Chiari, Caroline Zielinski, Yanlin Wang, Penghua Wang

## Abstract

Inflammasomes lead to activation of inflammatory caspases, which induce pyroptosis and an inflammatory immune response to control microbial infections. Inflammasomes are tightly regulated to avoid lethal sepsis and chronic autoimmune conditions. However, posttranslational regulation of inflammatory caspases remains poorly defined. We constructed 375 individual ubiquitin ligase knockout lines by CRISPR-Cas9, performed an unbiased screening, and identified Muscle Excess 3B (MEX3B), an RNA-binding protein and ubiquitin ligase, as a positive regulator of the caspase-4 inflammasome. Genetic depletion of MEX3B inhibited not only the caspase-4 but also NLRP3 and NLRC4 inflammasomes, regarding caspase activation, pyroptosis, and secretion of inflammasome-dependent cytokines, in human cells and murine primary macrophages. This MEX3B function required its RNA-binding, but not ubiquitin ligase activity. These results suggest that MEX3B is a pan-inflammasome regulator and a potential therapeutic target for inflammation.

## INTRODUCTION

Inflammasomes are large cytosolic multiprotein complexes formed in response to infections and cellular stresses, leading to proximity-induced auto-activation of inflammatory caspases (human caspase-1, -4, -5, murine caspase-1, -11), secretion of the interleukin (IL) −1β and −18 and pyroptosis, an inflammatory form of cell death (1). Canonically, in response to a pathogen ligand or cellular danger signal, a Pattern Recognition Receptor (PRR) oligomerizes into a platform, which recruits several caspase-1 molecules directly or via the adaptor, Apoptosis-associated Speck-like protein containing a Caspase recruitment domain (ASC). This allows for proximity-induced intermolecular auto-cleavage of caspase-1, which subsequently cleaves pro-IL-1β and pro-IL-18 into their biologically active, secreted forms. Active caspase-1 also cleaves gasdermin D (GSDMD) into an active N-terminal fragment that forms pores on the plasma membrane, leading to pyroptosis (1). Well-established inflammasomal PRRs include the Nucleotide-binding and Oligomerization Domain (NOD), Leucine-Rich Repeat (LRR)-containing Receptors (NLRP), Absent In Melanoma 2 (AIM2), Pyrin etc. However, the bacterial endotoxin lipopolysaccharide (LPS) can directly bind to human caspase-4/5 or rodent caspase-11, triggering their oligomerization and activation, known as the non-canonical inflammasome (2). These caspases can also directly cleave GSDMD and indirectly activate the NLRP3-ASC-caspase-1 inflammasome, causing cell death and maturation of the aforementioned proinflammatory cytokines, respectively. Mature IL-1β, −18 and other Damage Associated Molecular Patterns (DAMPs) are secreted via the GSDMD pores and further amplify downstream inflammation and immune cell activation, leading to pathogen clearance. However, aberrant activation of these inflammasome pathways may cause acute sepsis and chronic autoinflammatory / autoimmune conditions (3). Notably, sepsis constitutes ∼ 20% of all global deaths (4), outnumbering deaths from many common cancers and other diseases. The caspase-11 inflammasome is the predominant driver of LPS and polymicrobial sepsis and lethality in mice (5–7). Thus, inflammasomes are regulated at both the transcriptional and post-translational levels in cells to ensure proper functioning and signaling.

Ubiquitination is one of the most important post-translational modifications involved in many immune signaling pathways, including the NLRP3 inflammasome (8). For example, several ubiquitin E_3_ ligases are known to mediate degradative ubiquitination of NLRP3 (9–12), and a few promote NLRP3 assembly/activation via the non-degradative K63-linked polyubiquitination (13–16). However, there is a dearth of literature regarding the regulatory role of E_3_ ligases on the non-canonical inflammasomes (2). To date, only two E_3_ ligases have been shown to directly regulate the noncanonical inflammasome. Neural precursor cell Expressed Developmentally Down-regulated protein 4 (NEDD4) could induce K48-linked polyubiquitination and subsequent degradation of caspase-11 (17); and Synoviolin 1 (SYVN1) potentially promoting GSDMD activity via K27-linked polyubiquitination (18). To address the significant gap between ubiquitination and the caspase-4 inflammasome, we constructed 375 individual ubiquitin E_3_ ligase knockout lines by CRISPR-Cas9 (19) and performed an unbiased screening. This library represents almost all the currently known definite E3 ligases (total ∼377). Our screen identified 15 positive regulators of the caspase-4 inflammasome; one of the top hits was Muscle Excess 3B (MEX3B). The MEX3 family has 4 members, *i.e*., MEX3A-D, which are RNA-Binding Proteins (RBPs) and putative E_3_ ligases (20). However, their physiological functions remain poorly studied (21). Herein, we show that MEX3B promotes caspase and GSDMD activation, cell death, and secretion of inflammasome-dependent cytokines, upon activation of not only the caspase-4 but also NLRP3 and NLRC4 inflammasomes, in human cells and murine primary macrophages. Notably, this MEX3B function is dependent on its RNA-binding, not ubiquitin ligase activity. However, MEX3B is dispensable for the Toll-like receptor 4 (TLR4) and IFN-γ signaling. These results suggest that MEX3B is a pan-inflammasome regulator.

## RESULTS

### Identification of E_3_ ligases involved in the noncanonical inflammasome activation

The human genome encodes ∼617 ubiquitin E_3_ ligases and accessory proteins based on the presence of signature “catalytic” domains, as well as of domains distinctive of substrate recognition subunits of multi-subunit RING (Really Interesting New Gene) finger (RNF)-dependent E_3_ (22). However, only the HECT (Homologous to E6-associated protein C-Terminus), RING, and U-box proteins are considered definite E3 ligases (∼377 total). Using the E_3_ ligase CRISPR-Cas9 gRNA library from our previous work (19), we generated a library of 375 lentiviral particles from HEK293T, transduced A549 (a human lung cancer cell line) and selected stably transduced cells with puromycin (**Supplementary Fig. 1A**). We were unable to recover 20 knockout (KO) lines after antibiotic selection. The surviving E_3_ ligase KO cells were cultured overnight with human IFN-γ, transfected with LPS to activate the caspase-4 inflammasome (**Supplementary Fig. 1A**). We employed Lactate Dehydrogenase (LDH) release as a measurement of inflammasome activation/pyroptosis, and included both wild type (WT, scramble gRNA) and *CASP4^-/-^*cells as controls in each batch of screening.

A candidate E_3_ ligase was considered a positive or negative regulator if its KO led to a significant reduction [Log_2_(fold change over WT) ≦-1, Log_10_(p value) ≦-2] or increase [Log_2_(fold change over WT) ≧1, Log_10_(p value) ≦-2] in LDH release, respectively. With these criteria, we obtained 15 positive (blue dots) and 25 negative (red dots) regulators (**Supplementary Fig. 1B**). Of note, we recapitulated NEDD4 as a negative regulator of caspase-4 function from a recent study (17) (**Supplementary Fig. 1C**). For our study, we decided to focus primarily on the positive regulators as they can be the most therapeutically applicable. Of these, Tumor Necrosis Factor (TNF) Receptor Associated Factor 6 (TRAF6) (23, 24) and Ariadne RBR E3 Ubiquitin Protein Ligase 2 (ARIH2) (25) have been previously reported to regulate NLRP3 inflammasome function. To our knowledge, there is no literature illustrating a role for these 15 E3 ligases in the noncanonical inflammasome. We validated the results from the initial screen in our top positive regulators and all repeated well with a significant reduction in LDH release compared to WT cells (**Supplementary Fig. 1D**). Genetic deletion of MEX3B consistently led to the largest reduction in LDH release in A549 cells across numerous batches of experiments. For this reason, and because of the lack of research regarding the immunological function of MEX3B, we chose to explore MEX3B further to determine how it participates in the noncanonical inflammasome.

### MEX3B is critical for the noncanonical inflammasome activation

We next confirmed successful depletion of MEX3B and caspase-4 protein expression in *MEX3B^-/-^*and *CASP4*^-/-^ A549 cells, respectively (**Fig. 1A,B**). The GSDMD cleavage and pyroptosis in *MEX3B^-/-^* cells were reduced compared to WT, but to a less extent than *CASP4*^-/-^ (**Fig. 1B,C**). MEX3B belongs to the MEX3 family with four members, *i.e.,* MEX3A, -B -C, -D, which contain two evolutionary conserved RNA-binding K-homology (KH) domains and are potentially involved in mRNA decay (20,21). These MEX3-family members were included in our screen, with *MEX3A^-/-^* A549 cells exhibiting a slight decrease in LDH release while *MEX3C^-/-^* cells showing a slight increase (**Supplementary Fig. 1E**). *MEX3D^-/-^* cells did not survive puromycin selection. These results suggest a unique role for MEX3B in the caspase-4 inflammasome. Next, we reproduced the above findings in HeLa cells where the caspase-4 inflammasome signaling is more robust (**Fig. 1D-H**). Additionally, we noted that the secretion of inflammasome-dependent IL-18 and caspase-4 activation (p32 and active p20 fragments) were impaired in *MEX3B^-/-^* cells (**Fig. 1G,H**). Epichromosomal reconstitution of MEX3B expression to *MEX3B*^-/-^ HeLa cells restored normal caspase-4 inflammasome in terms of GSDMD cleavage and pyroptosis (**Fig. 1I,J**). We next explored the role of MEX3B in immune cells during inflammasome activation using THP-1 cells, a human monocytic cell line commonly used to study monocyte/macrophage function and inflammasomes. We generated polyclonal MEX3B knockout THP-1 cells (**Fig. 2A**) using CRISPR-Cas9, differentiated them into macrophages with phorbol 12-myristate 13-acetate (PMA), and transfected them with LPS to trigger the noncanonical inflammasome. We recapitulated the *MEX3B*^-/-^ phenotypes: reduced pyroptosis, GSDMD cleavage, and IL-1β and IL-18 release (**Fig. 2B-D**). Lastly, we confirmed these results using a human *MEX3B* siRNA in THP-1 cells (**Fig. 2E,F**). Taken together, these results provide solid evidence for a key role of MEX3B in optimal activation of the noncanonical inflammasome.

**Fig. 1.**
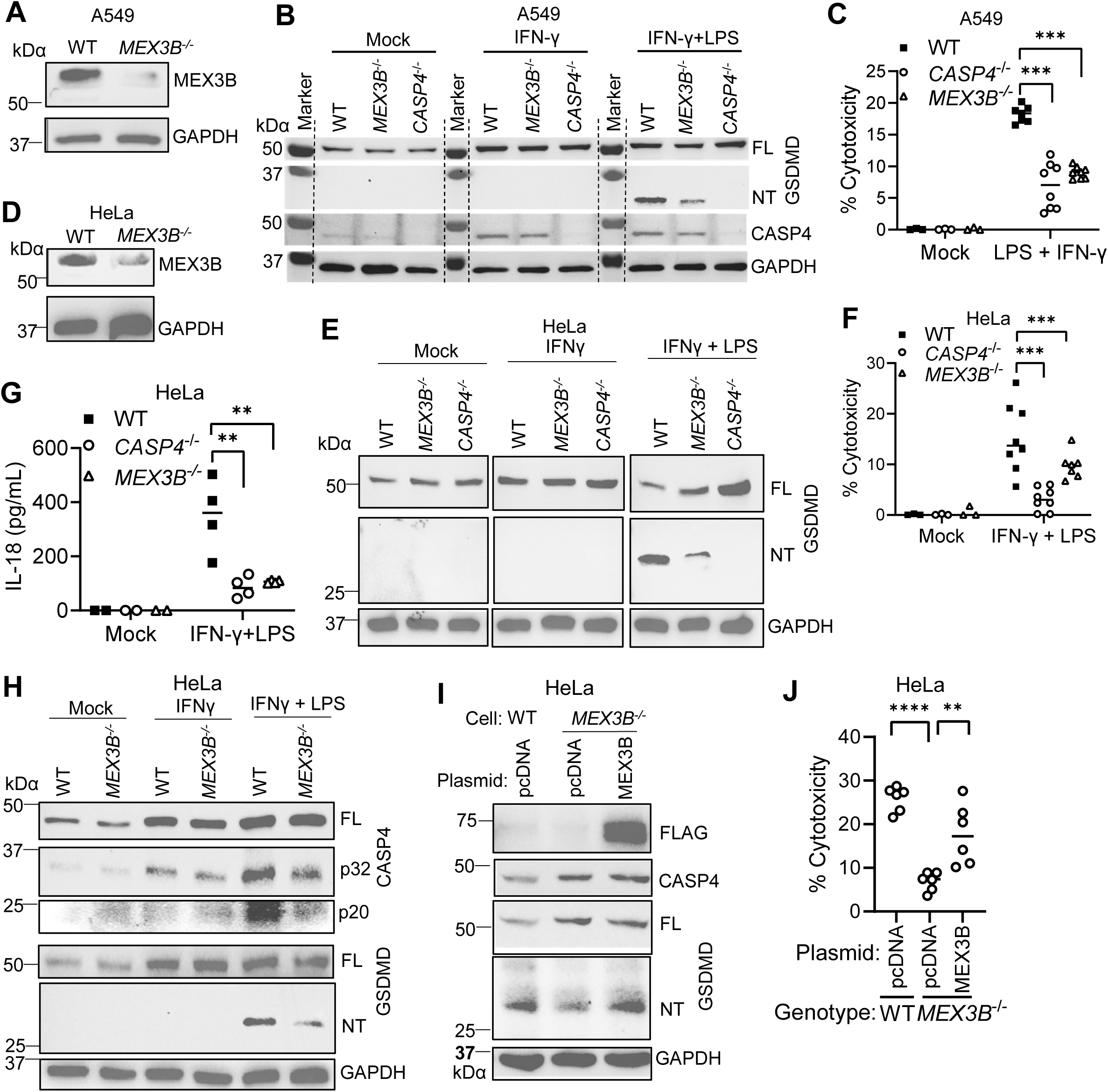
MEX3B is crucial for the caspase-4 inflammasome signaling in human epithelial cells. **A,B)** The immunoblots of **A)** MEX3B in A549 cells, **B)** caspase-4 and GSDMD in A549 cells primed with 100 ng/mL IFN-γ for 3 h, then transfected with 1 µg/mL LPS for 16 h. **C)** The percent cell death based on LDH release from A549 cells treated as in B). **D, E)** The immunoblots of **D)** MEX3B in HeLa cells, **E)** GSDMD in HeLa cells primed with IFN-γ for 16 h then transfected with 1 µg/mL of LPS for 6 h. **F)** The percent cell death based on LDH release and **G)** concentrations of secreted IL-18 from HeLa cells treated as in E). **H)** The immunoblots of caspase-4 and GSDMD in HeLa cells treated as in E). **H)** The immunoblots of caspase-4 and GSDMD, and **J)** percent cell death based on LDH release from HeLa cells transfected with plasmids for 24 h, followed by inflammasome stimulation as in E). FL, full-length, NT, N-terminus. kDα, molecular weight of protein ladders in kilodalton. C, F, G, J) Individual data plot with median, each symbol = one biological replicate, \*\**p*<0.01, *** *p*<0.001, **** *p*<0.0001 (ANOVA).

**Fig. 2.**
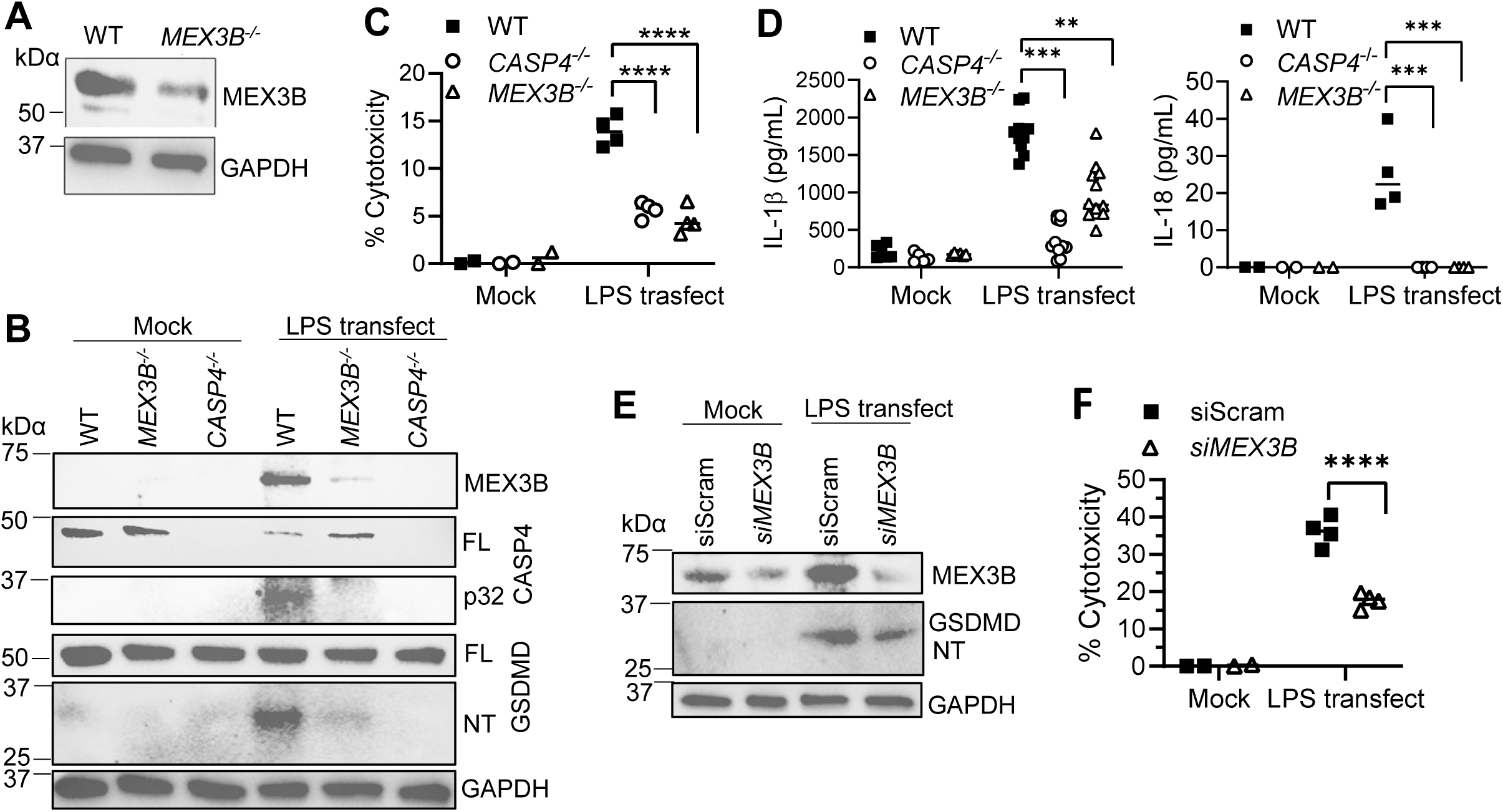
MEX3B is crucial for the caspase-4 inflammasome signaling in human macrophages. **A, B)** The immunoblots of **A)** MEX3B in THP-1 cells, **B)** caspase-4 and GSDMD in PMA-differentiated THP-1 macrophages transfected with 500 ng/mL LPS for 4 h. **C)** The percent cell death based on LDH release and **D)** concentrations of secreted IL-1β/IL-18 from THP-1 cells treated as in B). **E)** The immunoblots of indicated proteins and **F)** percent cell death from THP-1 macrophages transfected with siRNA for 48 h, then LPS for 4h. siScram, scrambled non-targeting control siRNA. FL, full-length; NT, N-terminus; kDα, molecular weight of protein ladders in kilodalton. C, D, F) Individual data plot with median, each symbol = one biological replicate, \*\**p*<0.01, *** *p*<0.001, **** *p*<0.0001 (ANOVA).

MEX3B is evolutionarily conserved, 96% identical between human and murine proteins. Next, we asked if MEX3B was also crucial for the noncanonical inflammasome in murine macrophages. We began with siRNA to transiently deplete *Mex3b* mRNA and protein expression in immortalized murine bone-marrow derived macrophages (iBMDMs) (**Fig. 3A,B**). The cleavage of both caspase-11 and GSDMD, and cell death and IL-1β release were reduced in cells Mex3b siRNA-transfected cells compared to those in scramble siRNA-treated cells; and the degree of reduction was correlated with *Mex3b* knockdown efficiency (**Fig. 3B-D**). Notably, these results were largely repeated in primary *Mex3b^-/-^* murine BMDMs (**Fig. 3E,F; Supplementary Fig. 2**).

**Fig. 3.**
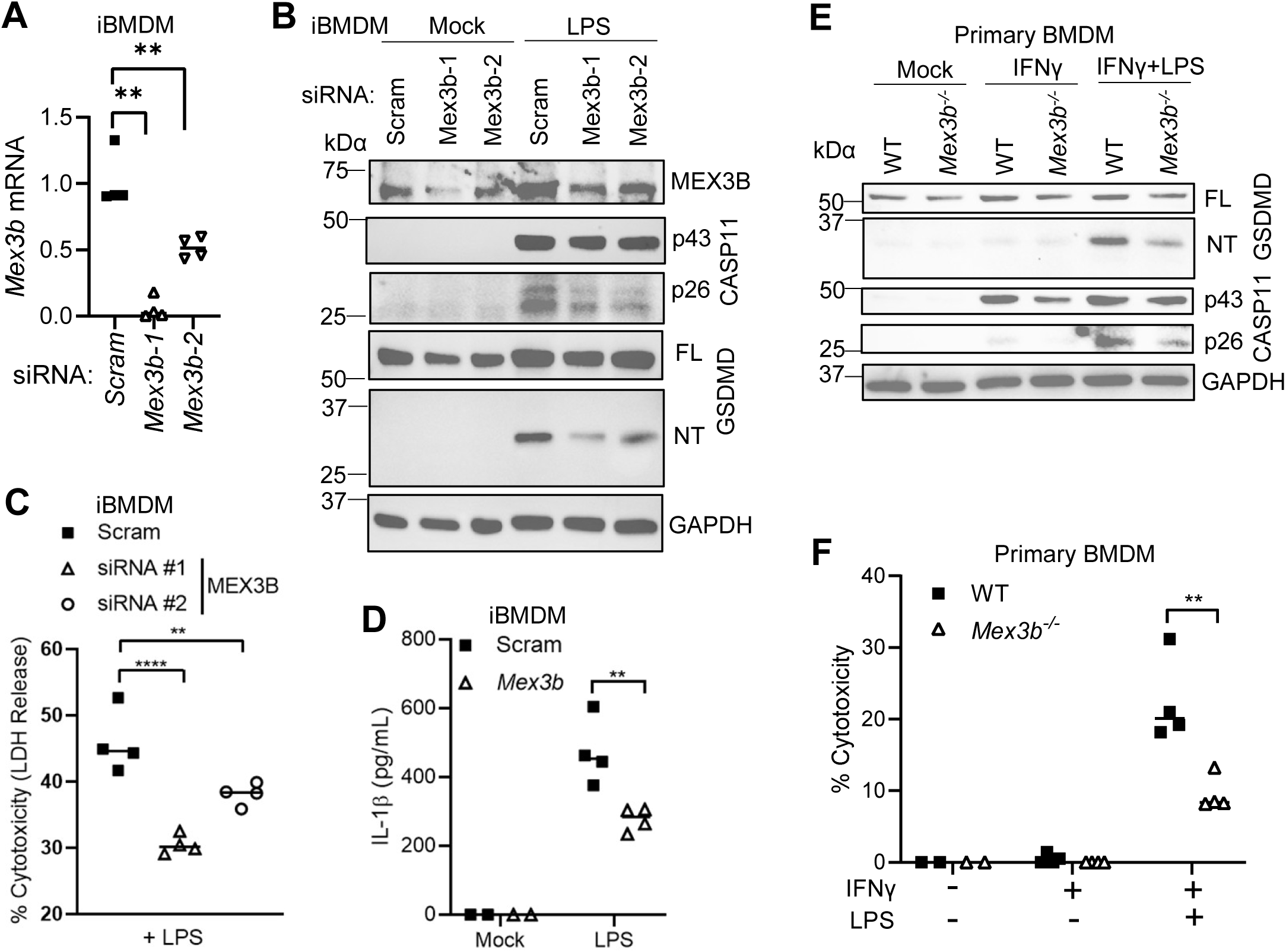
MEX3B is crucial for the caspase-11 inflammasome signaling in murine macrophages. **A)** qPCR quantification of *Mex3b* mRNA in immortalized bone-marrow derived macrophages (iBMDM) at 48 h after siRNA transfection. **B)** The immunoblots of indicated proteins in iBMDMs transfected with siRNA for 48 h, then 2.5 µg/mL of LPS for 6 h. **C)** The percent cell death based on LDH release, and **D**) IL-1β secretion, from iBMDMs treated as in **B**). **E)** The immunoblots of indicated proteins, and **F**) percent cell death, in primary BMDMs primed with 10 ng/mL of IFN-γ for 16 h then transfected with of 1 µg/mL LPS for 6h. A, C, D, F) Individual data plot, each symbol = one biological replicate, \*\**p*<0.01, **** *p*<0.0001 by ANOVA for A, C) or Student *t*-test for D, F).

### MEX3B is critical for canonical inflammasome activation

The aforementioned results have firmly established the conserved role for MEX3B in the non-canonical inflammasome. Next, we wanted to assess if MEX3B played a role in canonical inflammasomes. We first primed WT and *MEX3B^-/-^* THP-1 macrophages with LPS and activated NLRP3 with nigericin. We observed a notable decrease in the cleavage (activation) of caspase-1 and GSDMD, and cell death in *MEX3B^-/-^*cells, when compared to WT cells (**Fig. 4A,B**). These results were well reproduced in murine iBMDM and primary BMDMs (**Fig. 4C-F**). Moreover, flagellin-activated NLRC4 inflammasome signaling was inhibited in primary *Mex3b^-/-^* murine BMDMs (**Fig. 4G,H**). These results demonstrate that MEX3B is a positive pan-inflammasomal regulator targeting caspases.

**Fig. 4.**
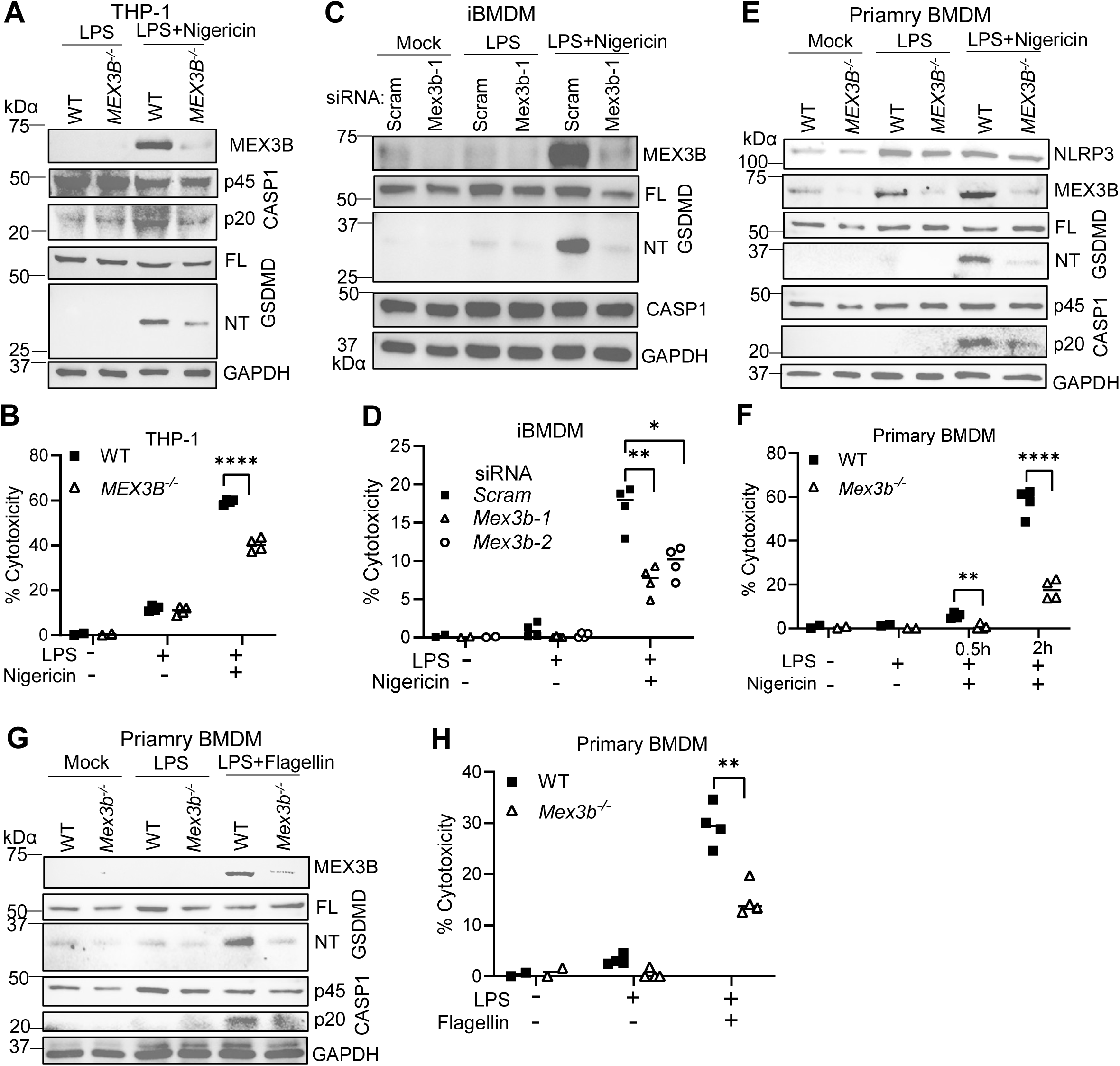
MEX3B is crucial for canonical inflammasome signaling. **A**) The immunoblots of indicated proteins and **B**) percent cell death based on LDH release, in human THP-1 macrophages primed with 100 ng/mL of LPS for 3 h then 10 µg/mL of nigericin for 2 h. **C**) The immunoblots of indicated proteins and **D**) percent cell death based on LDH release in immortalized murine BMDMs transfected with siRNA for 48 h, followed by stimulation with LPS/ nigericin as in **A**). **E**) The immunoblots of indicated proteins and **F**) percent cell death, in primary murine BMDM treated with LPS/ nigericin as in **A**). **G**) The immunoblots of indicated proteins and **H**) percent cell death, in primary murine BMDM treated with primed with 100 ng/mL of LPS for 3 h then transfected with 20 µg/mL of flagellin for 4 h. FL, full-length; NT, N-terminus; kDα, molecular weight of protein ladders in kilodalton. B, D, F, H) Individual data plot with median, n=4, \**p*<0.05, \*\**p*<0.01, *** *p*<0.001, **** *p*<0.0001 by Student *t*-test for B, F, H) or ANOVA for D).

### MEX3B is dispensable for the TLR4 and IFN-**γ** signaling

Robust inflammasome activation requires priming to upregulate expression of the inflammasome components such NLRP3, caspase-1/4 and GSDMD. In this study, we used IFN-γ or LPS to prime the caspase-4 or canonical inflammasome, via Toll-like receptor 4 (TLR4) or JAK-STAT1 signaling, respectively. Of note, a recent study revealed that MEX3B can act as a TLR3 co-receptor for viral RNA, playing an important role in innate antiviral responses (26). This prompted us to determine if MEX3B impacts the inflammasome-priming TLR4 or IFN-γ signaling pathways (27). Transcriptional upregulation of *Gsdmd*, *Casp11*, *Irf1/2* (transcriptional factors for *GSDMD*) (28) by IFN-γ was normal in *Mex3b^-/-^*, compared to WT BMDMs (**Supplementary Fig. 3A**). LPS-induced transcription of *Casp1/11*, *Tnf*, *Ifnb*, *Nlrp3* genes was intact too in *Mex3b^-/-^* BMDMs (**Supplementary Fig. 3B**). These results show that MEX3B is dispensable for these inflammasome-priming pathways.

### The RNA-binding feature of MEX3B is required for activating Caspase-4

The above dataset has firmly proven that loss of MEX3B function attenuates inflammasome signaling. We next asked if overexpression of MEX3B would reverse it. Indeed, either human or murine MEX3B overexpression increased GSDMD cleavage and cell death in HeLa cells (**Fig. 5A-D**). Notably, overexpression of MEX3B upregulated caspase-4 and GSDMD protein expression without/with IFN-γ priming (**Fig. 5A,C**). Together with the results from loss of MEX3B function, these results suggest that endogenous MEX3B is redundant for caspase-4/GSDMD expression. Indeed, overexpression of MEX3A also enhanced caspase-4 and GSDMD expression and cell death, while MEX3C failed to do so (**Supplemental Fig. 4A-C**).

**Fig. 5.**
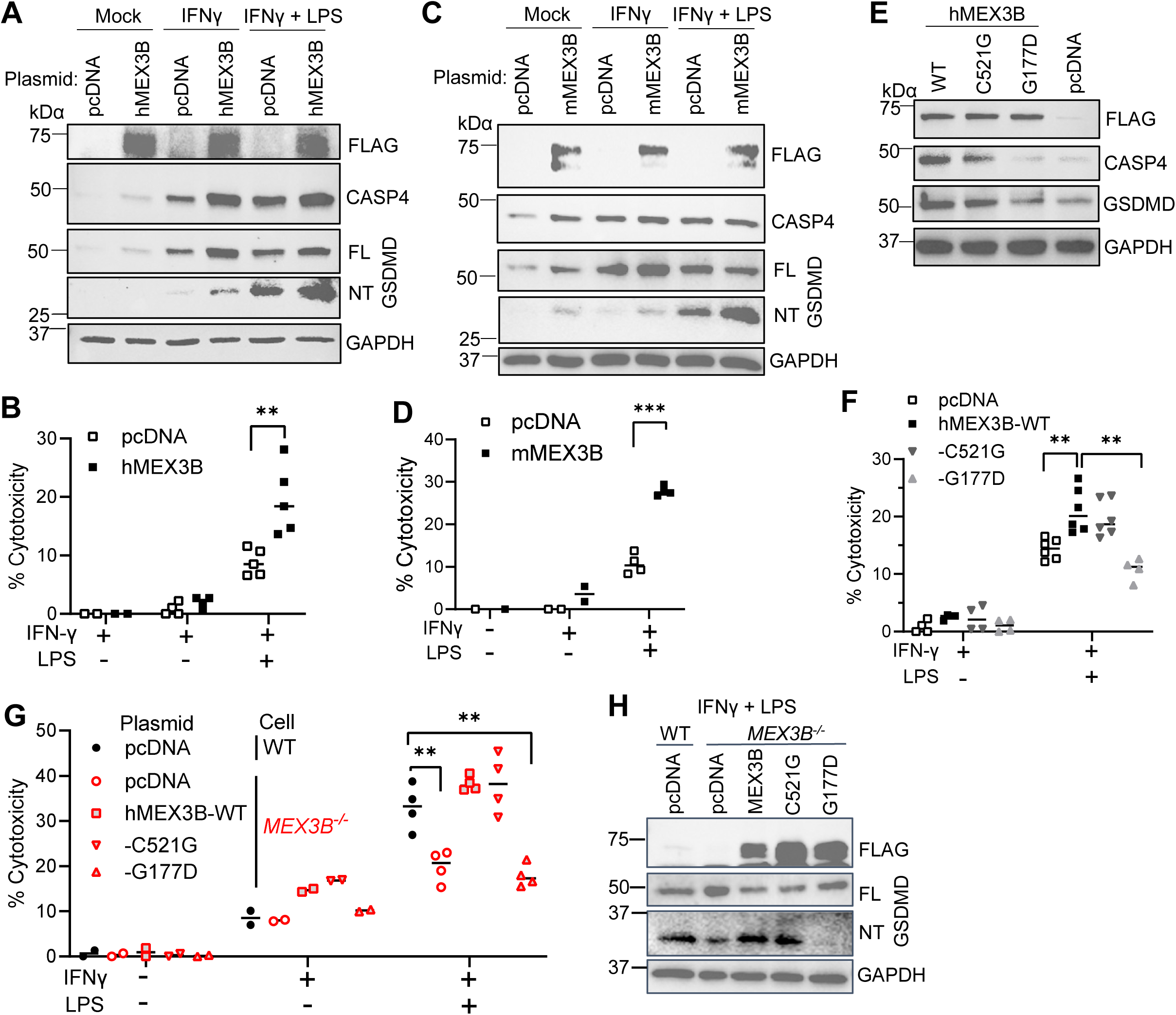
MEX3B requires its RNA-binding activity to promote inflammasome signaling. HeLa cells were transfected with a human (h)/mouse (m) MEX3B expression or empty vector (pcDNA) plasmid for 24 h, primed with 10 ng/mL of IFN-γ for 16 h, then transfected with 1 µg/mL of LPS for 6 h. WT, wild-type; C521G, mutation in ubiquitin ligase activity; G177D, mutation in RNA-binding activity. **A**, **C**, **E**, **H**) The immunoblots of indicated proteins. **B**, **D**, **F**, **G**) Percent cell death based on LDH release. FL, full-length; NT, N-terminus; kDα, molecular weight of protein ladders in kilodalton. B, D, F, G) Individual data plot with median, each symbol = one biological replicate, \*\**p*<0.01, *** *p*<0.001, by Student *t*-test for B, D) or ANOVA for F, H).

MEX3B has two RNA-binding KH domains at its N-terminus and one RING type ubiquitin E_3_ ligase domain at its C-terminus. A single mutation G177D in the second KH domain or C521G in the RING domain of MEX3B can abolish its RNA-binding or E_3_ ligase activity, respectively (20). To pinpoint the mechanism of MEX3B action on inflammasomes, we constructed G177D and C521G mutant human MEX3B expression plasmids by site-directed mutagenesis. Overexpression of the G177D mutant was unable to enhance GSDMD/caspase-4 protein levels and pyroptosis as WT MEX3B did (**Fig. 5E,F**). The C521G mutation reduced caspase-4 level by approximately 50%, however it did not influence pyroptosis, compared to WT MEX3B (**Fig. 5E,F; Supplemental Fig. 4D**). Moreover, while WT and the C521G mutant fully restored LPS-elicited pyroptosis and GSDMD cleavage to *MEX3B*^-/-^ cells, the G177D mutant was unable to do so (**Fig. 5G,H**). These results demonstrate that MEX3B requires its RNA-binding activity to promote caspase-4 inflammasome activation.

### Inflammasome activation increases MEX3B expression

We consistently noticed an increase in MEX3B protein expression during both the noncanonical (**Fig. 2B,E; Fig. 3B**) and canonical (**Fig. 4A,C, E, G**) inflammasome activation in macrophages, prompting us to investigate the role of inflammasome priming and activation on MEX3B expression. MEX3B protein expression in HeLa cells barely changed following IFN-γ priming alone, while increased substantially upon transfection with LPS (**Fig. 6A**). We next investigated the kinetics of MEX3B expression during caspase-4 inflammasome activation in THP-1 macrophages without priming. We noted that MEX3B protein level was upregulated synchronically with GSDMD cleavage, *i.e*., inflammasome activation in WT cells, and this upregulation was nearly ablated in *CASP4^-/-^* cells (**Fig. 6B**). MEX3B protein expression was induced in coincidence GSDMD cleavage in murine iBMDMs too (**Fig. 6C**). Consistent with MEX3B protein expression in HeLa cells, *MEX3B* mRNA expression increased modestly after IFN-γ priming, and much further after LPS transfection, and LPS transfection alone was sufficient to induce *MEX3B* mRNA in THP-1 cells (**Fig. 6D**). However, LPS priming alone reduced *Mex3b* mRNA expression by 4 times (**Fig. 6E**). Interestingly, in the MEX3 family, only *MEX3B* mRNA expression was responsive to IFN-γ/LPS priming or inflammasome activation (**Fig. 6D,E**), underlying the unique role of MEX3B in inflammasome activation.

**Fig. 6.**
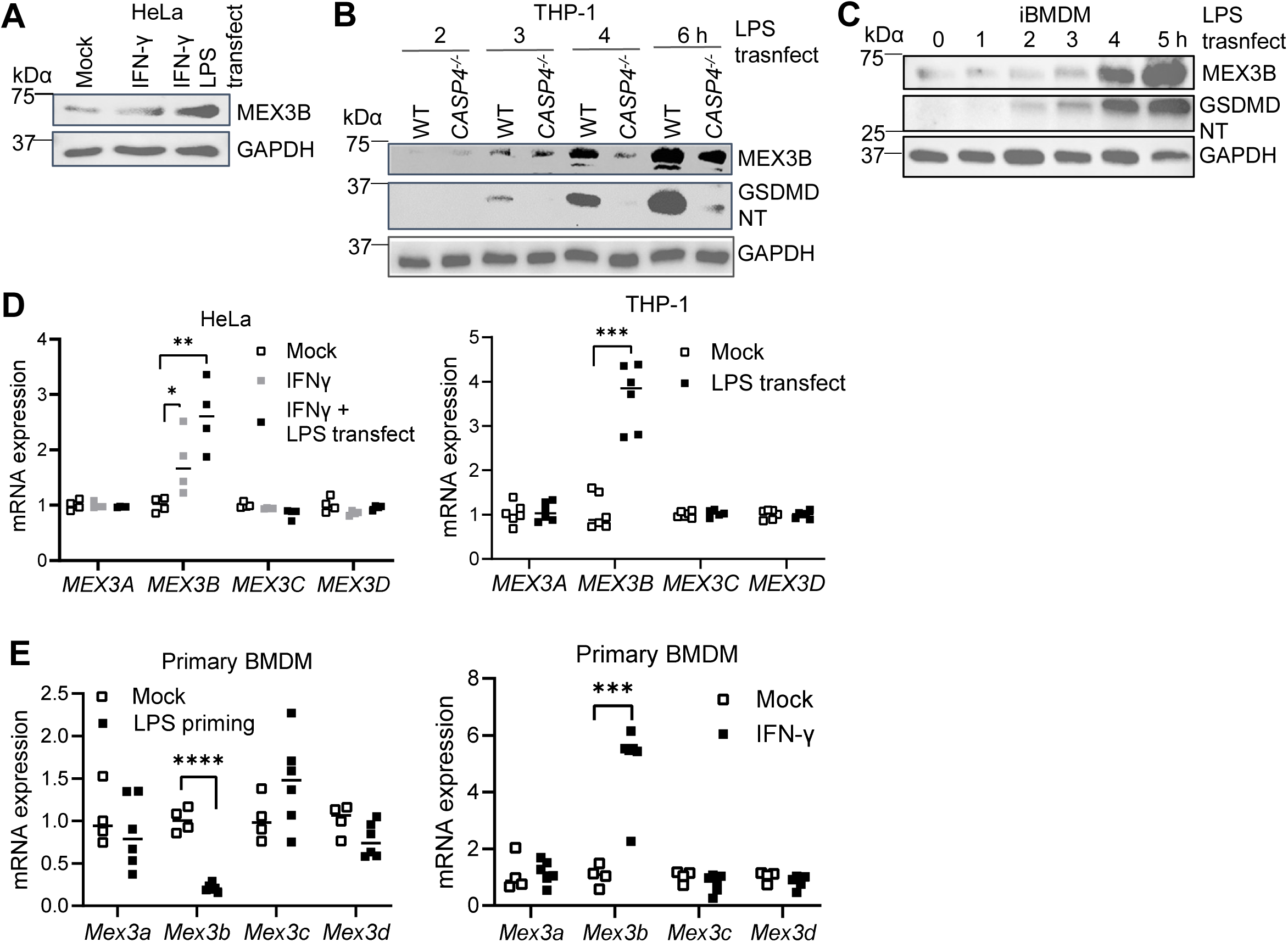
MEX3B expression is induced by inflammasome activation. **A-C)** The immunoblots of indicated proteins from **A)** HeLa cells primed with 10 ng/mL of human IFN-γ for 16 h then transfected 1 µg/mL with LPS for 6 h, **B)** PMA-differentiated THP-1 macrophages transfected with 500 ng/mL of LPS for 6 h, **C)** immortalized murine bone marrow-derived macrophages (iBMDMs) transfected with LPS. **D)** qPCR quantification of human *MEX3* mRNA levels in HeLa cells and THP-1 macrophages treated as in B, C). **E)** qPCR quantification of murine *Mex3* transcripts in primary BMDMs primed with 100 ng/mL of LPS for 3 h or IFN-γ for 16 h. NT, active N-terminal fragment; kDα, protein size markers in kilodalton. D, E) Individual data plot with median, each symbol = one biological replicate, \*\**p*<0.01, \*\*\**p*<0.001 by Student *t*-test.

## DISCUSSION

Inflammasomes are critical for controlling many microbial infections, however, aberrant activation of inflammasome signaling may lead to sepsis and autoinflammatory conditions. Cellular regulation of NLRP3 and ASC has been extensively studied, however, that of caspase-1/4 remains poorly explored (1). Here, we have identified MEX3B as a positive regulator of pan-inflammasomes.

MEX3B is a member of the MEX3 family with two evolutionarily conserved RNA-binding KH domains one ubiquitin ligase module (20). However, its physiological functions remain poorly studied (21). In *Caenorhabditis* (*C*.) *elegans,* null Mex-3 mutations (maternal effect) are embryonically lethal and render anterior blastomere (AB) of descendants to produce body wall muscle (should be made by posterior blastomere), hence the name Mex-3 for muscle excess 3 (29). In contrast to *C. elegans* Mex-3, mammalian MEX3 proteins contain a C-terminal RING (Really Interesting New Gene) E_3_ ligase domain (21). MEX3B E_3_ activity could mediate degradative ubiquitination of Suppressor of Zeste 12 homolog (SUZ12) (30) and Runt-Related Transcription Factor 3 (RUNX3) (31). Our results have demonstrated that the E_3_ ligase activity of MEX3B is largely dispensable, while its RNA-binding feature is required for, the caspase-4 inflammasome signaling (**Fig. 5G,H**). MEX3 proteins are known to bind to 3’-untranslated regions (UTR) of mRNAs to regulate their stability and translation. For instance, MEX3B could bind to the 3’-UTR of Human Leukocyte Antigen A (HLA-A) mRNA to destabilize it (31–33), degrades Transforming Growth Factor Beta Receptor III (TGFBR3) mRNA thus inhibiting collagen production(34). MEX3B could post-transcriptionally up-regulate IL-33 (35) and CXCL2 expression (36), thus increasing allergic airway inflammation. Our results show that genetic deletion of MEX3B did not obviously change the expression of inflammasome components including NLRP3, caspase-1/4 and GSDMD (**Figs.1-4**). However, transient overexpression of MEX3B modestly increased both caspase-4 and GSDMD transcript and protein levels, suggesting a redundant role of MEX3B in controlling caspase-4 and GSDMD expression. Indeed, overexpression of MEX3A also upregulated caspase-4/GSDMD expression (**Supplementary Fig. 4B, D**). Notably, among the MEX3 family, only MEX3B expression was upregulated by either IFN-γ priming or inflammasome signaling but reduced by LPS priming alone (**Fig. 6**). Moreover, at its C-terminus, MEX3B contains a serine-rich sequence (16 continuous serine residues), known as polyserine linker (PSL). PSLs serve as flexible, disordered spacers that connect different functional domains in modular proteins (37). These unique features in MEX3B might underlie its unique role in inflammasome signaling, among the MEX3 family.

In summary, we have revealed a critical role for MEX3B in activating both the canonical and non-canonical inflammasomes. The conserved RNA-binding domains of MEX3B may be therapeutically targeted to treat inflammasome-associated inflammatory diseases such as sepsis. Nonetheless, more work is needed to elucidate the molecular mechanism of MEX3B action.

## MATERIALS AND METHODS

### Reagents and antibodies

Antibodies for GAPDH (Cat# 60004-1-Ig, 1:2000) were purchased from Proteintech Group. Caspase-4 (Cat# 4450S, 1:1000), Gasdermin D (Cat# 39754S, 1:1000), Cleaved human-Gasdermin D (Asp275) (Cat# 36425S, 1:500), Cleaved mouse-Gasdermin D (Asp276) (Cat# 10137S, 1:1000), NLRP3 (Cat# 15101S, 1:1000), TBK1/NAK (Cat# 3504S, 1:1000), Phospho-TBK1/NAK (Ser172) (Cat # 5483S, 1:1000), p65 (Cat # 3034, 1:1000), Phospho-P65 (Cat # 3031S, 1:1000), and Actin (Cat# 4967S, 1:2000) antibodies were from Cell Signaling. Caspase-1 (Cat# ab179515, 1:1000), Caspase-11 (Cat# ab180673, 1:1000) antibodies were from Abcam. The Monoclonal Anti-FLAG M2 antibody (Cat# F1804, 1:2000), and lipopolysaccharides (E. coli O111:B4, Cat# L3024) were obtained from Sigma-Aldrich. Recombinant IFNγ protein of mouse (Cat# 485-MI-100) and human (Cat# 285-IF-100) were obtained from R&D systems. The MEX3B (Cat# PA5-72749, 1:1000) antibody, PureLink™ Genomic DNA Mini Kit (Cat# K182002), HiPure Plasmid Filter Midiprep Kit (Cat# K210014), and Quick Plasmid Miniprep Kit (Cat# K210010) were from Thermo Fisher. The TransIT-X2 Dynamic Delivery System (Cat# MIR6005) was from Mirus Bio LLC.

### Murine model

All the animal procedures were approved by the Institutional Animal Care and Use Committee at UConn Health adhering to federal and state laws. Sperm from C57BL/6NCrl-Mex3btm1b(KOMP)Wtsi/Ph mice (RRID:IMSR_EM:11612) was purchased via the European Mouse Mutant Archive (EMMA) from the Institute of Molecular Genetics (Czech Republic). *In vitro* fertilization (IVF) was performed by the Center for Comparative Medicine at UConn Health. Heterozygous mice were then bred to generate *Mex3b^-/-^* mice, where the critical exon was flanked by loxP sites, and subsequent Cre expression excised this critical sequence resulting in a knockout reporter allele [Gt(ROSA)26Sortm1(ACTB-cre,-EGFP)Ics (MGI:5285392)]. These mice were housed in the specific pathogen-free animal facility at UConn Health. All mice were housed at an ambient temperature of approximately 24 °C, a humidity of 40∼60%, and a light/dark cycle of 12 h. Genotyping was performed with genomic DNA and Choice Taq Blue Mastermix (Denville Scientific, Cat# CB4065-8) under the following PCR: 95 °C for 5 min, 34 cycles of 95 °C for 30 sec, 62 °C for 30 s, 72 °C for 1 min, and then 72 °C for 5 min, 4 °C to stop. The genotyping primers for wild type *Mex3b* were Forward: 5‘-ctttgggagggggcagt-3’, Reverse: 5‘-ctggctccttgtctcgg-3’. For the mutant *Mex3b*, the primers were Forward: 5‘-acggtttccatatggggatt-3’, Reverse: 5‘-tctttgccataaggccatactaa-3’. The PCR reaction resulted in products of 871 bp (WT) and/or 698 bp (transgene).

### Differentiation of primary bone marrow cells into macrophages

Bone marrow cells were isolated and differentiated into macrophages (BMDM) using the established methods^19^. Briefly, bone marrow cells were obtained from mouse hind legs by centrifugation at 8000rpm for 1 min and differentiated in L929-conditioned medium (RPMI 1640, 20% FBS, 30% L929 culture medium, 1x antibiotics/antimycotics) in a 10-cm Petri dish at 37 °C, 5% CO2 for 5-7 days, with a change of medium every 2 days. Attached BMDMs were dislodged by incubating in ice-cold PBS for 30 min followed by pipetting and counted for plating in cell culture plates. The BMDMs were seeded and treated in plating medium (RPMI 1640, 10% FBS, 5% L929-conditioned medium, 1x antibiotics / antimycotics).

### Cell culture

HeLa (Adenocarcinoma epithelial cell, Cat# CCL-2), HEK293T cells (human embryonic kidney, Cat# CRL-3216), A549 (human lung carcinoma epithelial cell, Cat# CCL-185), THP-1 (human acute monocytic leukemia, Cat# TIB-202) and L929 cells (mouse fibroblast cells, Cat# CCL-1) were purchased from the American Type Culture Collection (ATCC) (Manassas, VA 20110, USA). HEK293T, HeLa and L929 cells were grown in Dulbecco’s modified Eagle’s medium (DMEM, Corning, Cat#10-013-CV) supplemented with 10% fetal bovine serum (FBS, Gibco™, Cat#A52567-01) and 1% antibiotics/antimycotics (ThermoFisher Scientific, USA). A549 and THP-1 cells were cultured in Roswell Park Memorial Institute (RPMI) 1640 (Corning, Cat# 10-040-CV) supplemented with 10% fetal bovine serum and 1% antibiotics/antimycotics. We obtained our immortalized bone-marrow derived macrophages (iBMDMs) as a gift from Dr. Vijay Rathinam. These cells were cultured in DMEM with 10% fetal bovine serum and 1% antibiotics/antimycotics. All cells were grown in an incubator set to 37_°C and 5% CO_2_.

### Generation of gene knockout cell lines using CRISPR-Cas9

Gene knockout in human and mouse cell lines by CRISPR-Cas9 were performed based on our previous studies (Reference). Briefly, pre-designed, gene unique guide (g) RNAs (Integrated DNA Technologies, USA) (Sequences in Supplemental Table 1) were subcloned into a lentiCRISPR-V2 vector (Reference) and correct insertion was confirmed by DNA sequencing. To generate lentiviral particles, each gRNA vector was transfected into HEK293T cells with the packaging plasmids pCMV-VSV-G (Reference) and psPAX2 (#12259, from the Didier Trono lab via Addgene, USA). Half of the cell culture medium was replaced with fresh medium at 24 h and viral particles were collected at 48-72 h after transfection. The cell culture supernatant (viral particles) was cleared by brief centrifugation at 2000 rpm for 5 min. Target cells at ∼50% confluence in a 12-well plate were transduced with 1 mL of each lentiviral preparation and 8 μg/mL polybrene for 24 h and selected with 1_µg/mL of puromycin in growth media for 6 days. The cells were then cultured in fresh complete RPMI or DMEM for experiments or frozen for stock. The knockout efficiency of the top candidates was tested by the T7EI assay for gene editing efficiency (Reference) or by protein expression via western blot.

### siRNA transfection

siRNA oligo duplexes for mouse-Mex3a (Cat # SR416592), mouse-Mex3b (Cat # SR417646), and human-MEX3B (Cat # SR313397) were obtained from Origene and prepared following the manufacturer’s instructions. Initially, three individual siRNAs for each protein were provided and tested at a concentration of 20 nM for their ability to knock down mRNA of these transcripts via RT-qPCR. The individual siRNA that displayed the best knockdown efficiency was used in future experiments. After siRNA transfection via the TransIT-X2 Dynamic Delivery System, cells were cultured for ∼60 hours before any additional experiments began. siRNA controls (20 nM) were included in each batch of experiments.

### Induction of inflammasomes

A549, HeLa, THP-1, L929, and iBMDM cells were cultured as described above. A549 cells were primed with 100 ng/mL of human IFN-γ for 3 h and then transfected with 1 μg/mL of LPS (E. coli O111:B4) for 18 h using TransIT-X2 Dynamic Delivery System. HeLa cells followed a similar methodology, except were primed with 10ng/mL of human IFN-γ for 18 h and then transfected with 1 μg/mL of LPS for 6 h. THP-1 cells were first differentiated into macrophages with 10ng/mL of phorbol myristate acetate (PMA) for 72 h followed by a 24 h rest period. Afterwards, these macrophages were transfected with 500 ng/mL of LPS for 4 h. L929 cells were primed with 50ng/mL of mouse IFN-γ for 18 h and then transfected with 5 μg/mL of LPS for 6 h. Lastly, iBMDM cells were transfected with 2.5 μg/mL of LPS for 5 h. For plasmid overexpression-induced inflammasome responses and complementation (i.e. rescue) studies, HeLa cells were first transfected with the specific plasmid for 24 h and then primed with IFN-γ and transfected with LPS as described earlier. In all cases, after induction, cell lysates and supernatants were collected for immunoblotting or assays for cytokines and lactate dehydrogenase (LDH) release, respectively.

Induction of canonical inflammasome in macrophages began first with LPS priming (100 ng/mL for 3 h), followed by addition of 10 μg/mL of nigericin (NLRP3 inflammasome) (Cat# N7143-5MG, Sigma-Aldrich) to the cell culture media for 1-3 h, or transfection with 20 µg/mL of flagellin (NLRC4 inflammasome) for 4 h. After induction, cell lysates and supernatants were collected for immunoblotting or assays for cytokines and lactate dehydrogenase (LDH) release, respectively.

### Extraction of total RNA and RT-qPCR

Cells were collected in 350□µL of lysis buffer (RNApure Tissue & Cell Kit, CoWin Biosciences, USA), containing 1% β-mercaptoethanol. Total cellular RNA was extracted according to the product manual. Reverse transcription of RNA into complementary DNA (cDNA) was performed using the PrimeScript™ RT reagent Kit (Takara Bio, Inc, Cat# RR037A). Quantitative PCR (qPCR) was performed with gene-specific primers and iTaq Universal SYBR Green Supermix (BioRad, Cat# 1725124). Results were calculated using the –ΔΔCt method and a housekeeping gene (ACTB) as an internal control. The qPCR primers are listed in Supplemental Table 2.

### Cell death and cytokine assays by ELISA

Cell death was analyzed by measuring LDH release into the cell culture medium with the Cytotoxicity Detection KitPLUS (LDH) kit (Roche, Cat# 04744926001) according to the manufacturer’s protocol. The release of IL-1β or IL-18 in cell culture supernatants were quantified by: ELISA MAX™ Deluxe Set human IL-1β (BioLegend, Cat# 437015) and mouse IL-1β DuoSet ELISA kit (R&D, Cat# DY401-05), while human IL-18 was quantified via the DuoSet ELISA IL-18 Kit (R&D, Cat# DY318-05).

### Immunoblotting

Immunoblots were performed using a standard method based on our previous studies^3^. In brief, protein samples were diluted with 2x or 4x Laemmli protein sample buffer (Bio-Rad, Cat# #1610747) followed by 100 °C boiling for 10 min, and loaded to 15 well 4%-15% Mini-PROTEAN® TGXTM Precast protein gels (Bio-Rad) for standard sodium dodecyl sulfate-polyacrylamide gel electrophoresis (SDS-PAGE), Western blotting, and an enhanced chemiluminescent (ECL) substrate (ThermoFisher, Cat# 32106) or an ultra-sensitive Lumigen ECL Ultra substrate for low-femtogram-level detection (LUMIGEN, Cat# TMA-100) were applied. Primary antibodies used were diluted in a 5% fat-free milk solution. An anti-rabbit IgG, HRP-linked secondary antibody (Cell Signaling Technology, Cat#7074) and anti-mouse IgG, HRP-linked secondary antibody (Cat#7076) were used at a dilution of 1:5000 in 5% fat-free milk. Protein band detection was performed using a ChemiDoc MP Imaging System (Bio-Rad). The band density was quantified by Image J.

### Semi-denaturing detergent agarose gel electrophoresis (SDD-AGE)

Cell lysates from primary mouse macrophages treated as previously described were combined with 4x SDD-AGE loading buffer and run on a 1.5% agarose gel with 0.1% SDS for 1h, 100V @□4°C. Afterwards, the proteins from the gel were transferred onto a 0.45µm nitrocellulose membrane for immunoblotting. Antibody staining and imaging were performed as described in the immunoblotting methods section. The NLRP3 antibody (Cat# AG-20B-0014-C100, 1:1000) was purchased from AdipoGen.

### Plasmid construction/point mutagenesis

Human FLAG-MEX3B was constructed by inserting the human MEX3B open reading frame (ORF) (NCBI accession: NM_032246.6.) into the pcDNA3.1-FLAG vector (Reference) using standard PCR amplification and cloning techniques. The pcDNA-FLAG-MEX3B plasmids were transformed into *Escherichia coli* DH5α (ThermoFisher Scientific, Cat # 18265017) and plated on LB Agar containing 100□µg ml−1 ampicillin and inverted overnight at 37° C. Antibiotic-resistant colonies were picked and grown in LB media at 37° C overnight by shaking (100□rpm). Plasmid DNA was extracted using PureLink Quick Plasmid Miniprep Kit (ThermoFisher Scientific, Cat # K210011). Sanger sequencing was used to verify MEX3B insertion into plasmids. All other FLAG-tagged plasmids (e.g. FLAG-MEX3A) were cloned following a similar protocol. To create the point mutants of MEX3B, we used the QuikChange II Site-Directed Mutagenesis Kit (Cat # 200523, Agilent) and followed the manufacturers protocol. Our Human FLAG-MEX3B served as a template to clone these mutants (-C215G, -G177D). PCR primers for these mutants were purchased from Integrated DNA Technologies. For primer sequences, see Supplementary Table 3).

### Graphing and statistical analyses

Graphpad Prism was used for graphing and statistical analyses. The sample sizes for animal experiments were estimated according to our prior experience in similar experiments. Survival curves were analyzed using a log-rank (Mantel–Cox) test. A two-tailed, unpaired Student’s *t* test or non–parametric/parametric Mann–Whitney U test was applied to a data set statistical analysis depending on its data distribution. The one-way or two-way ANOVA test was applied to simultaneously comparing multiple groups or with multiple time points. Data were presented in mean ± S.E.M, *p* values ≤0.05 were considered significant. The sample sizes, statistical tests, and *p* values were specified in figure legends.

## Data availability

The datasets generated during and/or analyzed during the current study are in the supplemental materials. Source data are provided with this paper.

## Acknowledgements

This work was supported by the National Institutes of Health of Unites States grant R01AI187147 to P.W. and a UConn Health startup fund to P.W.

## Author contributions

J.G.C. performed most experimental procedures, acquired and analyzed the data. T.G., D.Y, C.C., H.C., N.K.T. assisted J.G.C. with some experimental procedures and / or provided technical support. Y.W. provided access to essential instruments for data collection. J.G.C. and P.W. wrote the manuscript. P.W. conceived and supervised the study. All the authors reviewed and/or modified the manuscript.

## REFERENCES

1. D. Zheng, T. Liwinski, E. Elinav, Inflammasome activation and regulation: toward a better understanding of complex mechanisms. Cell Discov 6, 36 (2020).

2. J. Cahoon, D. Yang, P. Wang, The noncanonical inflammasome in health and disease. Infectious Medicine 1, 208–216 (2022).

3. N. Kelley, D. Jeltema, Y. Duan, Y. He, The NLRP3 Inflammasome: An Overview of Mechanisms of Activation and Regulation. Int J Mol Sci 20 (2019).

4. K. E. Rudd et al., Global, regional, and national sepsis incidence and mortality, 1990-2017: analysis for the Global Burden of Disease Study. Lancet 395, 200–211 (2020).

5. S. M. Man et al., Differential roles of caspase-1 and caspase-11 in infection and inflammation. Sci Rep 7, 45126 (2017).

6. M. Deng et al., The Endotoxin Delivery Protein HMGB1 Mediates Caspase-11-Dependent Lethality in Sepsis. Immunity 49, 740–753 e747 (2018).

7. A. J. Russo et al., Intracellular immune sensing promotes inflammation via gasdermin D-driven release of a lectin alarmin. Nat Immunol 22, 154–165 (2021).

8. J. Xu, G. Nunez, The NLRP3 inflammasome: activation and regulation. Trends Biochem Sci 10.1016/j.tibs.2022.10.002 (2022).

9. S. Han et al., Lipopolysaccharide Primes the NALP3 Inflammasome by Inhibiting Its Ubiquitination and Degradation Mediated by the SCFFBXL2 E3 Ligase. J Biol Chem 290, 18124–18133 (2015).

10. H. Song et al., The E3 ubiquitin ligase TRIM31 attenuates NLRP3 inflammasome activation by promoting proteasomal degradation of NLRP3. Nat Commun 7, 13727 (2016).

11. J. Tang et al., Sequential ubiquitination of NLRP3 by RNF125 and Cbl-b limits inflammasome activation and endotoxemia. J Exp Med 217 (2020).

12. D. Wang et al., YAP promotes the activation of NLRP3 inflammasome via blocking K27-linked polyubiquitination of NLRP3. Nat Commun 12, 2674 (2021).

13. F. Humphries et al., The E3 ubiquitin ligase Pellino2 mediates priming of the NLRP3 inflammasome. Nat Commun 9, 1560 (2018).

14. K. Labbe, C. R. McIntire, K. Doiron, P. M. Leblanc, M. Saleh, Cellular inhibitors of apoptosis proteins cIAP1 and cIAP2 are required for efficient caspase-1 activation by the inflammasome. Immunity 35, 897–907 (2011).

15. J. Ni et al., Ubc13 Promotes K63-Linked Polyubiquitination of NLRP3 to Activate Inflammasome. J Immunol 206, 2376–2385 (2021).

16. L. Zhang et al., Peli1 facilitates NLRP3 inflammasome activation by mediating ASC ubiquitination. Cell Rep 37, 109904 (2021).

17. Q. Liu, S. Zhang, Z. Sun, X. Guo, H. Zhou, E3 ubiquitin ligase Nedd4 is a key negative regulator for non-canonical inflammasome activation. Cell Death Differ 26, 2386–2399 (2019).

18. Y. Shi et al., E3 ubiquitin ligase SYVN1 is a key positive regulator for GSDMD-mediated pyroptosis. Cell Death Dis 13, 106 (2022).

19. D. Yang et al., UBR5 promotes antiviral immunity by disengaging the transcriptional brake on RIG-I like receptors. Nat Commun 15, 780 (2024).

20. K. Buchet-Poyau et al., Identification and characterization of human Mex-3 proteins, a novel family of evolutionarily conserved RNA-binding proteins differentially localized to processing bodies. Nucleic Acids Res 35, 1289–1300 (2007).

21. S. Jasinski-Bergner, A. Steven, B. Seliger, The Role of the RNA-Binding Protein Family MEX-3 in Tumorigenesis. Int J Mol Sci 21 (2020).

22. W. Li et al., Genome-wide and functional annotation of human E3 ubiquitin ligases identifies MULAN, a mitochondrial E3 that regulates the organelle’s dynamics and signaling. PLoS One 3, e1487 (2008).

23. H. W. Chiu, C. H. Chen, J. N. Chang, C. H. Chen, Y. H. Hsu, Far-infrared promotes burn wound healing by suppressing NLRP3 inflammasome caused by enhanced autophagy. J Mol Med (Berl) 94, 809–819 (2016).

24. Y. Xing et al., Cutting Edge: TRAF6 Mediates TLR/IL-1R Signaling-Induced Nontranscriptional Priming of the NLRP3 Inflammasome. J Immunol 199, 1561–1566 (2017).

25. A. Kawashima et al., ARIH2 Ubiquitinates NLRP3 and Negatively Regulates NLRP3 Inflammasome Activation in Macrophages. J Immunol 199, 3614–3622 (2017).

26. Y. Yang et al., The RNA-binding protein Mex3B is a coreceptor of Toll-like receptor 3 in innate antiviral response. Cell Res 26, 288–303 (2016).

27. N. Kopitar-Jerala, The Role of Interferons in Inflammation and Inflammasome Activation. Front Immunol 8, 873 (2017).

28. N. Kayagaki et al., IRF2 transcriptionally induces GSDMD expression for pyroptosis. Sci Signal 12 (2019).

29. B. W. Draper, C. C. Mello, B. Bowerman, J. Hardin, J. R. Priess, MEX-3 is a KH domain protein that regulates blastomere identity in early C. elegans embryos. Cell 87, 205–216 (1996).

30. H. Zhang et al., RNA helicase DEAD box protein 5 regulates Polycomb repressive complex 2/Hox transcript antisense intergenic RNA function in hepatitis B virus infection and hepatocarcinogenesis. Hepatology 64, 1033–1048 (2016).

31. M. Xue et al., HOTAIR induces the ubiquitination of Runx3 by interacting with Mex3b and enhances the invasion of gastric cancer cells. Gastric Cancer 21, 756–764 (2018).

32. L. Huang et al., The RNA-binding Protein MEX3B Mediates Resistance to Cancer Immunotherapy by Downregulating HLA-A Expression. Clin Cancer Res 24, 3366–3376 (2018).

33. K. Yang et al., Molecular mechanism of specific HLA-A mRNA recognition by the RNA-binding-protein hMEX3B to promote tumor immune escape. Commun Biol 7, 158 (2024).

34. J. X. Liu, et al., MEX3B inhibits collagen production in eosinophilic nasal polyps by downregulating epithelial cell TGFBR3 mRNA stability. JCI Insight 8 (2023).

35. Y. Yamazumi et al., The RNA Binding Protein Mex-3B Is Required for IL-33 Induction in the Development of Allergic Airway Inflammation. Cell Rep 16, 2456–2471 (2016).

36. Y. Yamazumi et al., The RNA-binding protein Mex-3B plays critical roles in the development of steroid-resistant neutrophilic airway inflammation. Biochem Biophys Res Commun 519, 220–226 (2019).

37. H. Shen et al., Deletion of the linker connecting the catalytic and cellulose-binding domains of endoglucanase A (CenA) of Cellulomonas fimi alters its conformation and catalytic activity. J Biol Chem 266, 11335–11340 (1991).

