## Supplementary Figures 1-4 for "MEX3B is a positive pan-inflammasome regulator"

This supplementary material contains Supplementary Figs.1-4 and legends.

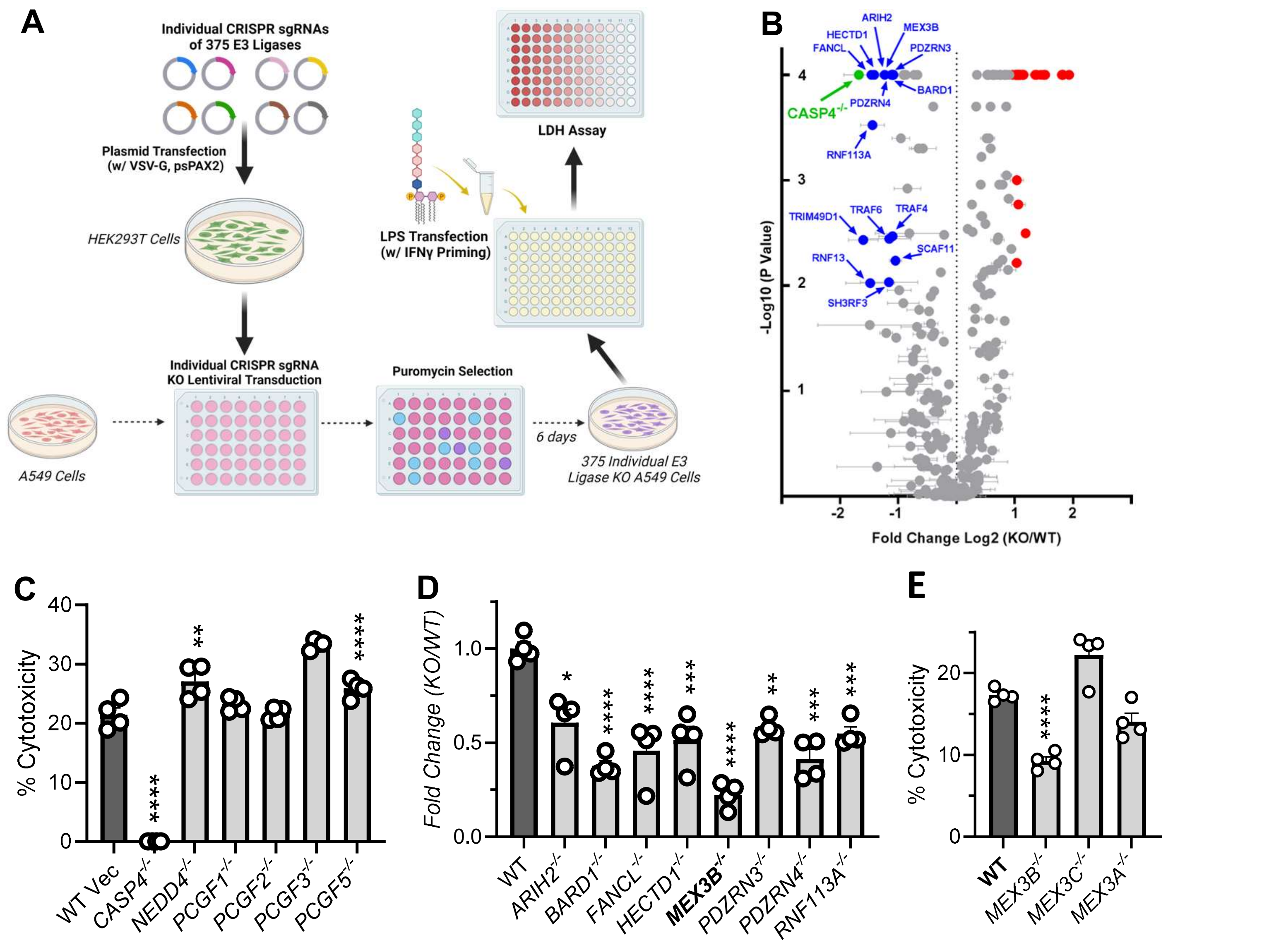


**Supplementary Fig.1. Identification of E3 ligases involved in the non-canonical inflammasome signaling. A)** The workflow of screening with 375 individual E3 ligase knockout (KO) A549 lines. **B**) The volcano plot of fold changes in the extracellular LDH (cell death readout) concentrations, **C, E**) Percent cell death, and **D**) fold change in LDH, of individual KOs over wild type (WT). Bar, mean ± S.E.M., N=4, **p*<0.05, ***p*<0.01, ****p*<0.001, *****p*<0.0001 (ANOVA).

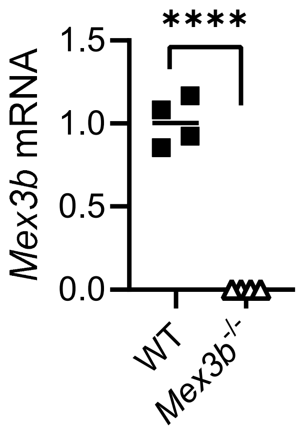


**Supplementary Fig.2. Validation of *Mex3b*$^{-/-}$ mice.** Shown is the individual data plot of qRT-PCR quantification of *Mex3b* mRNA expression in primary murine bone marrow-derived macrophages (BMDMs). **** $p<0.0001$ (Student *t*-test), n=4.

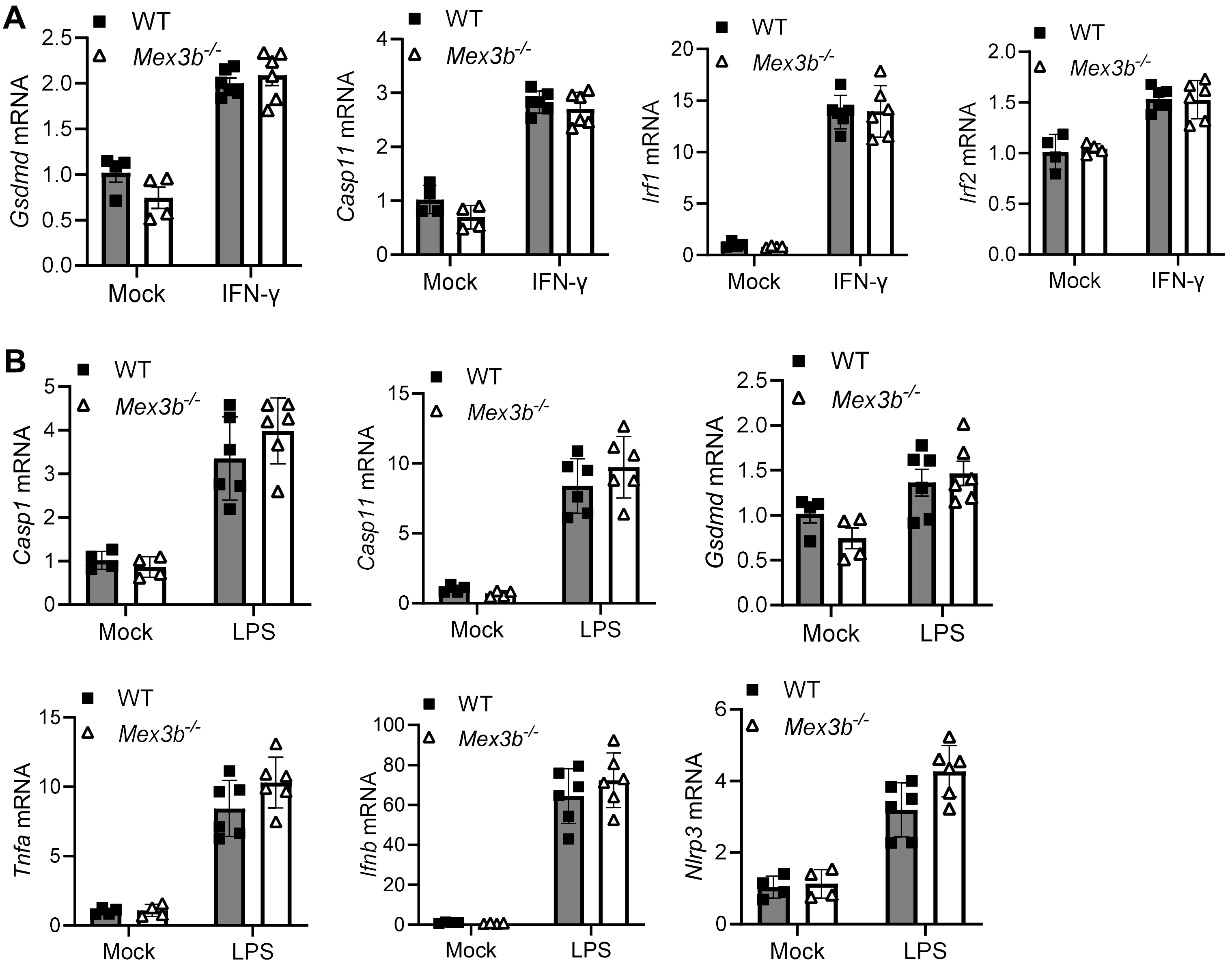


**Supplementary Fig.3. MEX3B is dispensable for TLR4 and IFN-γ signaling.** qRT-PCR quantification of indicated gene mRNA expression in primary murine bone marrow-derived macrophages (BMDMs) primed with **A)** 10 ng/mL of IFN-γ for 16 h, or **B)** 100 ng/mL of LPS for 3 h. Bar, mean ± S.D.; N=4-6.

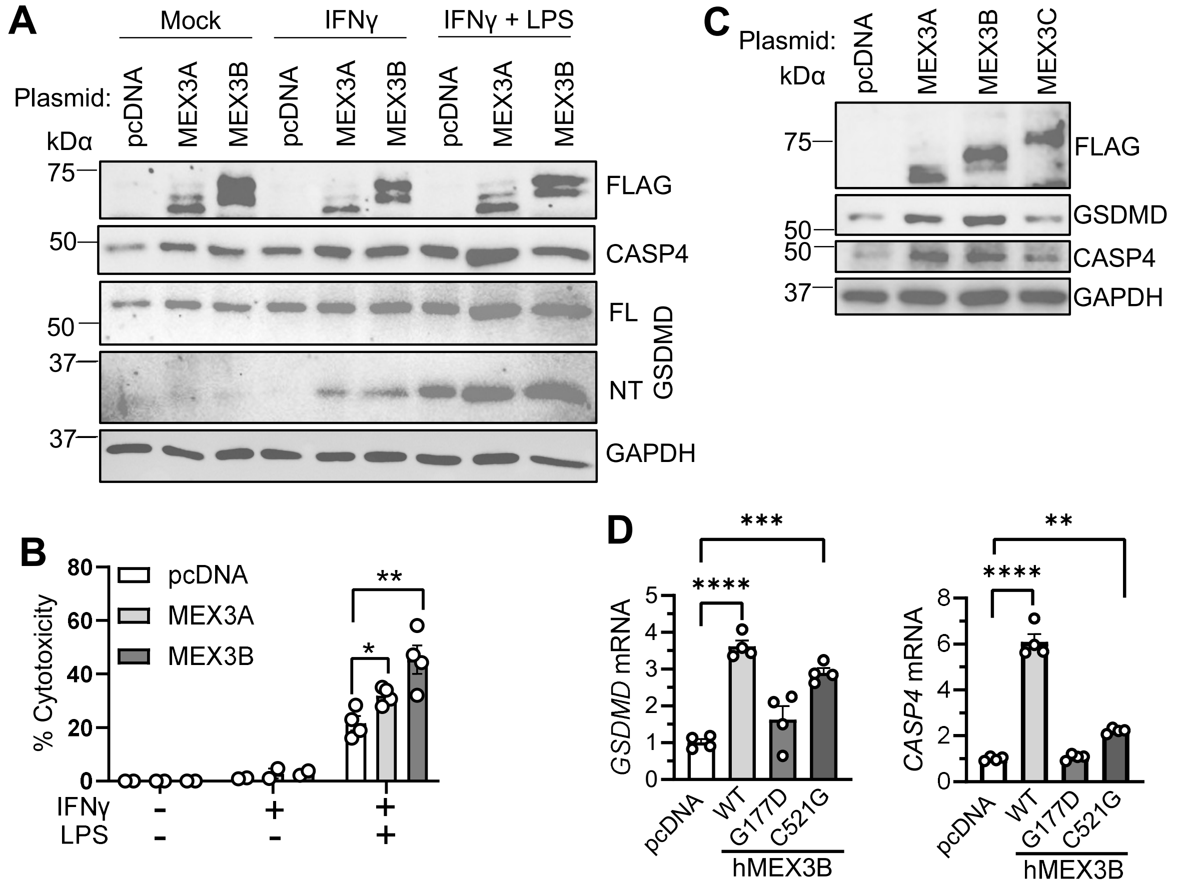


**Supplementary Fig.4. Overexpression of MEX3B increases caspase-4 and GSDMD expression. A, B)** HeLa cells were transfected with human MEX3 expression or empty vector (pcDNA) plasmids for 24 h, primed with 10 ng/mL of IFN-γ for 16 h, then transfected with 1 μg/mL of LPS for 6 h. **A)** The immunoblots of indicated proteins, **B)** percent cell death based on LDH release. **C**) The immunoblots of indicated proteins, and **D)** qPCR quantification of indicated mRNA levels in HeLa cells transfected with human MEX3 plasmids for 24 h. WT, wild-type; C521G, mutation in ubiquitin ligase activity; G177D, mutation in RNA-binding activity. Bar, mean ± S.E.M., N=4, *$p<0.05$, **$p<0.01$, *** $p<0.001$, **** $p<0.0001$ by ANOVA.
